# Deep Video Analysis for Bacteria Genotype Prediction

**DOI:** 10.1101/2024.09.16.613253

**Authors:** Ali Dabouei, Ishan Mishra, Kuwar Kapur, Chengzhi Cao, Andrew A. Bridges, Min Xu

## Abstract

Genetic modification of microbes is central to many biotechnology fields, such as industrial microbiology, bioproduction, and drug discovery. Understanding how specific genetic modifications influence observable bacterial behaviors is crucial for advancing these fields. In this study, we propose a supervised model to classify bacteria harboring single gene modifications to draw connections between phenotype and genotype. In particular, we demonstrate that the spatiotemporal patterns of *Vibrio cholerae* growth, recorded in terms of low-resolution bright-field microscopy videos, are highly predictive of the genotype class. Additionally, we introduce a weakly supervised approach to identify key moments in culture growth that significantly contribute to prediction accuracy. By focusing on the temporal expressions of bacterial behavior, our findings offer valuable insights into the underlying mechanisms and developmental stages by which specific genes control observable phenotypes. This research opens new avenues for automating the analysis of phenotypes, with potential applications for drug discovery, disease management, etc. Furthermore, this work highlights the potential of using machine learning techniques to explore the functional roles of specific genes using a low-resolution light microscope.

## 1 Introduction

The presence of bacteria, among other microorganisms, is vital for Earth’s ecosystem and life support systems. They contribute to essential ecological processes, such as nutrient cycling, aiding plant growth, and the decomposition of organic materials, ultimately ensuring the health of ecosystems. In the case of human health, bacteria have a large impact, both positive and negative. Beneficial bacteria in the human body aid in digestion, produce essential vitamins, and contribute to a well-functioning immune system. On the other hand, in many cases, bacteria are notorious for causing infectious diseases and dysbiosis [1, 2]. Many bacteria exhibit rapid growth rates and undergo rapid mutagenesis and horizontal gene transfer enabling them to overcome challenges such as antibiotic treatment. For instance, pathogens such as *Vibrio cholerae* and *Escherichia coli* can double approximately every 20 minutes under rich growth conditions, and through time have developed increasing resistance to commonly used antibiotics. Controlling bacterial growth, and the behaviors that bacteria exhibit in the infection process, are key topics of research that can contribute to drug design [3]. Leblanc and Charles [4] highlight the challenges of working with bacterial cells due to their metabolism, genetic stability, and the toxicity of the products being developed. The study of how particular genes influence bacterial growth and behaviors is crucial for defining new targets for drug discovery [5–7]. Controlling bacterial lifestyles is a fundamental challenge in microbiology that can be aided by modern computational approaches, and have applications in various fields. Understanding and manipulating the timing of bacterial behaviors is crucial in processes such as fermentation for the production of biofuels and pharmaceuticals [8]. The ability to design new strategies for controlling bacterial behavior via computational approaches helps understand possible optimizations to larger biological problems.

Defining genes that are responsible for particular bacterial behaviors involves generating specific gene modifications by experimental scientists to influence the way bacteria grow, interact, and respond to their environment. This process often includes the deletion or disruption of particular genes, and the subsequent impact on bacterial behavior is analyzed. For example, disrupting genes associated with antibiotic resistance can enhance the effectiveness of antibacterial drugs. Understanding how gene modifications influence bacterial behavior is essential for advancing applications such as drug development and environmental remediation. However, challenges lie in the nature of these genetic modifications. The complexity of each modified gene’s effect on bacterial behavior is a significant hurdle. Furthermore, when multiple genes are altered simultaneously, predicting the outcome becomes complex [9], as the cumulative impact may not be in line with the outcome of modifying each individual gene.

In this work, we aimed to develop a model for analyzing and classifying the behavior of the global pathogen, *Vibrio cholerae*, harboring single-gene deletions. To this end, we performed low-magnification brightfield timelapse microscopy of culture growth for the wildtype parent strain and 9 additional strains harboring single-gene modifications known to impact bacterial behavior. Our goal was to draw connections between phenotype and genotype. To this end, we implement a video analysis approach to predict the disrupted gene based on the culture growth videos.

In addition to classifying mutant phenotypes, we present a novel weakly supervised approach that identifies the saliency (contribution) of video frames in predicting the gene classes given the weak supervision of the video class during the training. The integration of saliency scores offers a novel perspective on video data analysis. By assigning significance scores to specific frames in a bacteria’s lifecycle, we analyze and understand patterns in the modified bacteria’s behavior.

Additionally, we introduce GeneMod, the dataset used for training and evaluating our model, consisting of 849×25 CLIP [10]feature embeddings of *Vibrio cholerae* culture growth.

## 2 Related Works

### 2.1 Gene Modification

Genetic modification has revolutionized various fields and provided findings to various challenges such as vaccine development, environmental remediation, and food production [11–13]. In particular, it has played a key role in the development of vaccines [14–16] and has enabled the sustainable production of genetically modified crops on a large scale helping food production [13, 17–19]. Additionally, genetic modification finds numerous applications in environmental remediation [20–22]. For instance, it has been used in the development of bioengineered microorganisms for oil spill cleanups [23]. Furthermore, genetic modification increases the efficiency of wastewater treatment processes [24, 25], reducing the environmental impact of industrial and municipal wastewater. Genetic modification has emerged as a powerful tool across various fields and promising a more sustainable future.

### 2.2 Sequential Data Analysis

The analysis of sequential data is essential in numerous domains, including natural language processing, time series forecasting, and biological data analysis. Traditional methods such as Hidden Markov Models (HMM) [26] and simple Recurrent Neural Networks (RNN) [27] were adopted for sequence classification and prediction. Although these techniques showed promise, they struggled to capture long-range dependencies in sequences effectively. Long Short-Term Memory (LSTM) networks were introduced by Hochreiter & Schmidhuber [28], offering improved memory capabilities and the ability to alleviate vanishing gradient problem [29]. LSTM became generally accepted for tasks involving sequential data owing to their ability to retain information over longer sequences. Following LSTM, Gated Recurrent Unit (GRU) [30] networks emerged as a simpler alternative, offering comparable performance with fewer parameters [31]. GRU achieved this by combining the forget and input gates of LSTM into a single update gate, reducing computational complexity while retaining the ability to capture temporal dependencies effectively. However, the most significant leap in sequential data analysis came with the development of Transformer architectures [32]. Transformers introduced a self-attention mechanism, allowing the model to weigh the significance of different elements in the input sequence dynamically.

Sequential modelling techniques have proven to be invaluable in biology [33–35]. HMM [36–38] and RNN [39] have been instrumental in the interpretation of proteomics and genomics. LSTM networks [40–42] and GRU [41, 43] have shown promising results in drug discovery and metabolomics. Transformers have revolutionized the exploration of longer sequences [44, 45], providing essential insights in evolutionary biology. Furthermore, the usage of these sequential models is not restricted to those mentioned and they have been used across the entire spectrum of biological research [46–48].

### 2.3 Saliency Detection

Targeting the identification of pivotal features within input samples, saliency detection emerges as a crucial technique with broad applications across across interdisciplinary fields [49, 50]. It has been used in the medical domain in the space of medical imaging [51–53]. It has also proven to be crucial in designing autonomous vehicles controlling their ability to safely perceive and navigate complex environments [54–56]. This has subsequently aided remote sensing applications [57, 58], by enabling the identification of features and patterns in satellite or aerial imagery for environmental monitoring and disaster relief. In addition, saliency recognition in robotics [59–61] has aided in object recognition, obstacle avoidance, and scene understanding, contributing to the further development of autonomous robotic systems.

In our work, we utilize saliency prediction to pinpoint salient frames, particularly to capture significant moments in the bacteria’s life cycle, using a weakly supervised approach. ‘Weakly supervised methods’ do not imply direct monitoring of saliency scores; We predict saliency indirectly using the monitoring provided as class labels. This approach involves identifying frames in input videos that have the greatest influence on the prediction outcome. Although this is extensively researched in the field of computer vision [62–65], the application of it in the context of bacteria growth is not explored.

## 3 Materials and Methods

Here, we present our approach for classifying bacterial growth videos to identify specific genetic modifications. The organization of the subsections are as follows: We start with subsection Video acquisition, where we discuss the process of acquisition of the modified video samples, this is followed by subsection Preprocessing, preprocessing is described that covers a series of essential steps to enhance the video data for subsequent analysis. Moving forward to the subsection Feature Extraction, we describe the importance of feature extraction, highlighting the critical role of data preparation and feature extraction. In the subsection Temporal Analysis, we discuss temporal analysis using a Transformer-based model to process the bacteria growth video. In the subsection Weakly-supervised Saliency Detection, we introduce a weakly supervised saliency detection module that allows us to identify salient time stamps of the bacteria’s life cycle, and in the subsection Classification, we present the final module that performs the classification to estimate the gene modification class.

### 3.1 Video acquisition

Brightfield timelapse microscopy of *V. cholerae* growth (parent strain WT O1 El Tor biotype C6706str2) was performed as described previously [66].Selected mutant strains were chosen due to their known role in regulating particular bacterial behaviors such as biofilm formation and motility. Briefly, cultures of the indicated strains were pre-grown overnight in 96-well microtiter dishes (Corning) in 200 uL of LB media. Overnight cultures were subsequently diluted 200,000-fold into M9 media containing 0.5% dextrose and 0.5% casamino acids in new microtiter dishes. Subcultured strains were grown statically at 30 degrees Celsius in a BioSpa incubator and were robotically transferred for imaging at one-hour intervals for 24 hours total. Brightfield microscopy was performed with fixed imaging conditions, a Biotek Cytation 1 instrument equipped with a 10x objective lens.

### 3.2 Preprocessing

Initially, a local contrast normalization [67–69] operation is applied with a block radius of 100 in both the x and y directions and a standard deviation of 3. Local Contrast Normalization is a type of normalization that works by enhancing an image’s features while reducing variability between different parts of the image, making it easier to distinguish different features when analyzing them later. This step aims to improve the visibility of structures in the video, contributing to better feature extraction. The data is converted to an 8-bit format for standardization. With that, a Gaussian blur with a sigma value of 2 is employed to reduce noise and smooth the image stack. A few examples of certain video frames before and after preprocessing can be seen in figure 2.

### 3.3 Feature Extraction

Data preparation and feature engineering are important processes that greatly affect video analysis performance. The primary purpose of these processes is to include noise reduction, feature selection, selecting important representation features, and transforming high-dimension features into the subspace domain without losing valuable information. For this, we use the feature representation learned via Contrastive Language-Image Pre-training (CLIP) [10]. CLIP is a foundational neural network trained on millions of images, which is used to understand visual concepts with the supervision of textual descriptions. CLIP’s unique fusion of vision and language understanding has proven to be highly versatile and adaptable to multiple domains [70, 71]. Moreover, the versatility and adaptability of CLIP provide meaningful embeddings for the frames by effectively using vision understanding. We use CLIP with ViT-b/32, ViT-b/16, ResNet101, and ResNet50 as the base models, to extract those high-dimension features to be used for training our model. Formally, we denote the CLIP feature extractor as `CLIP` : ℝ^*h×w×c*^ *→*ℝ^*d*^, where *h, w*, and *c* are the height, width, and the number of channels in the input frames, and *d* is the cardinality of the output space.

Let 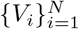 denote the set N bacteria growth videos, preprocessed using the criteria mentioned in Section Preprocessing. For the *j*^*th*^ frame of the *i*^*th*^ video, *v*_*i,j*_ *∈ V*_*i*_, we extract features as *x*_*i,j*_ = CLIP(*v*_*i,j*_). Then, we concatenate the computed CLIP features to come up with the set of features for each video that, 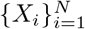 denote the set of CLIP embeddings for all the extracted frames.

### 3.4 Temporal Analysis

The video data recorded from bacterial growth is a sequence of *n*_*f*_ frames capturing the specific timestamp in the bacteria’s life cycle. In the previous section, we extracted features from the frames of the video using the CLIP backbone. The CLIP captures spatial information in each of the frames. Processing these data requires a second model to capture the temporal dependency of the features extracted from the frames. Recurrent Neural Networks capable of analyzing sequential data are the best choice for this aim since they can draw connections between different time samples in the data and combine them to compute the final prediction. This is a very important step in the processing since different bacteria with different gene modifications may express similar behavior at different timestamps of their lifecycle. Hence, the temporal analysis enables us to differentiate between these cases.

Transformers [32] are the most capable recurrent networks and have revolutionized many fields including NLP, time series analysis, computer vision, etc. The Transformer architecture is known for capturing long-range information in sequential data [72]. The core of our model is the transformer layer, which is based on the Transformer architecture. It employs multi-head self-attention mechanisms to analyze the input data’s contextual relationships. Additionally, it uses a feedforward neural network with a hidden dimension. We use the transformer model with *n*_*l*_ encoder layers to process temporal information and predict the class associated with each of the video inputs. Formally, the transformer encoder in our model is denoted as 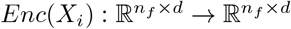, where *n*_*f*_ is the number of frames in each video and *d* is the dimension of CLIP model’s embedding.

### 3.5 Weakly-supervised Saliency Detection

Our analysis shows that we can achieve notable performance in predicting the classes of gene modifications from observing the bacteria growth videos. However, it is unclear which frames or moments in the bacteria’s life cycle are more informative about their modified gene. Hence, in this section, We have explored the development of a crucial component, Temporal Saliency aimed at assessing the significance of individual frames within bacteria growth videos. This temporal component is designed to give us extra insights into the life cycle of bacteria. It contributes to the decision-making process, by enhancing our understanding through analyzing multiple frames captured during their life cycle.

Specifically, the Saliency is used to focus on the relevant frames of the input data [73]. It produces a saliency map that highlights the most important or salient region in a specific frame. The main purpose of a saliency map is to identify areas that are considered significant by a computational model. In Figure 5 we can see the comparison of the average saliency map for each class computed with the help of ViT-b/32 and ResNet50 backbone of CLIP. From the figure, we can see how the graph dynamically responds to changing frames, this representation signifies the importance of specific frames across multiple frames for a given class.

This is integrated into the architecture to handle Temporal attention. It takes the same input as the Transformer and generates a saliency map of the same dimensions using a linear network. Formally, the Weakly-supervised Saliency Detection in our model is denoted as 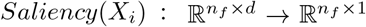, where *n*_*f*_ is the number of frames in each video and *d* is the dimension of CLIP model’s embedding. Notably, we adjust the size of the saliency map to align with the dimensions of the output from our transformer model. The resulting saliency map is then utilized to regulate the output of the transformer encoder. The outcome represents the aggregate representation that emphasizes the important frames of a bacteria’s lifecycle at distinct timestamps. Before passing this aggregate representation to the Classifier Classification, a mean operation is applied, ensuring a refined input for further stages of the model. This whole approach allows us to prioritize relevant frames, thereby enhancing the model’s ability to recognize bacterial behavior throughout its life cycle.

### 3.6 Classification

The Classifier is a custom neural network that takes input as the mean aggregation of the output from the transformer encoder multiplied by the saliency map which we discussed in Weakly-supervised Saliency Detection. The central objective of our custom neural network is to classify between a set of 10 classes, within the target frame. It consists of a fully connected layer (FC) with an output dimension of 10, for 10 distinct classes. Fig 1 provides an overview of our approach where an aggregate sequence representation is passed through a fully connected layer, followed by a softmax layer, to output probabilities for each of the 10 respective classes.

**Figure 1:**
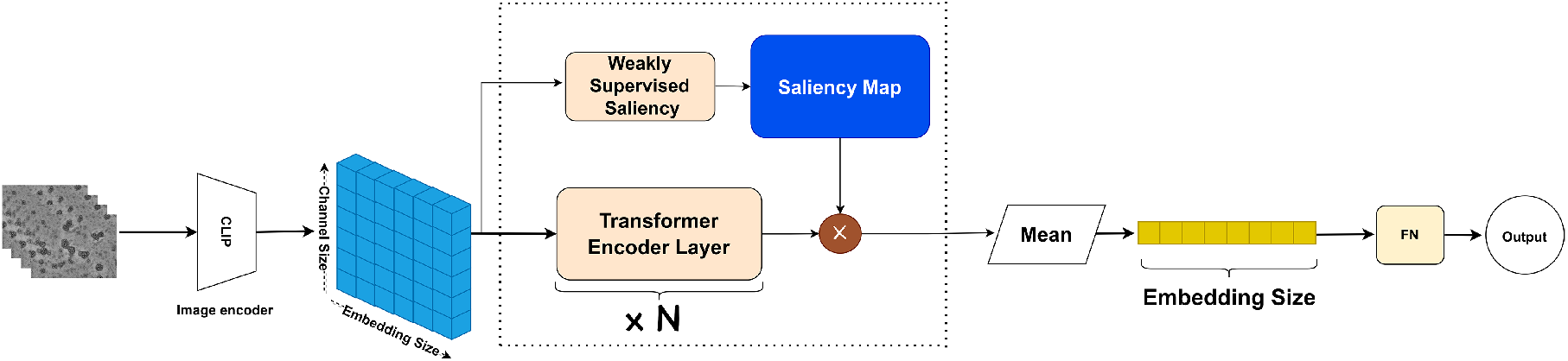
Illustration of the proposed model architecture for bacteria growth classification. The schematic diagram integrates a weakly supervised saliency detection model, estimating the contribution of each frame in the input video to the final prediction.

**Figure 2:**
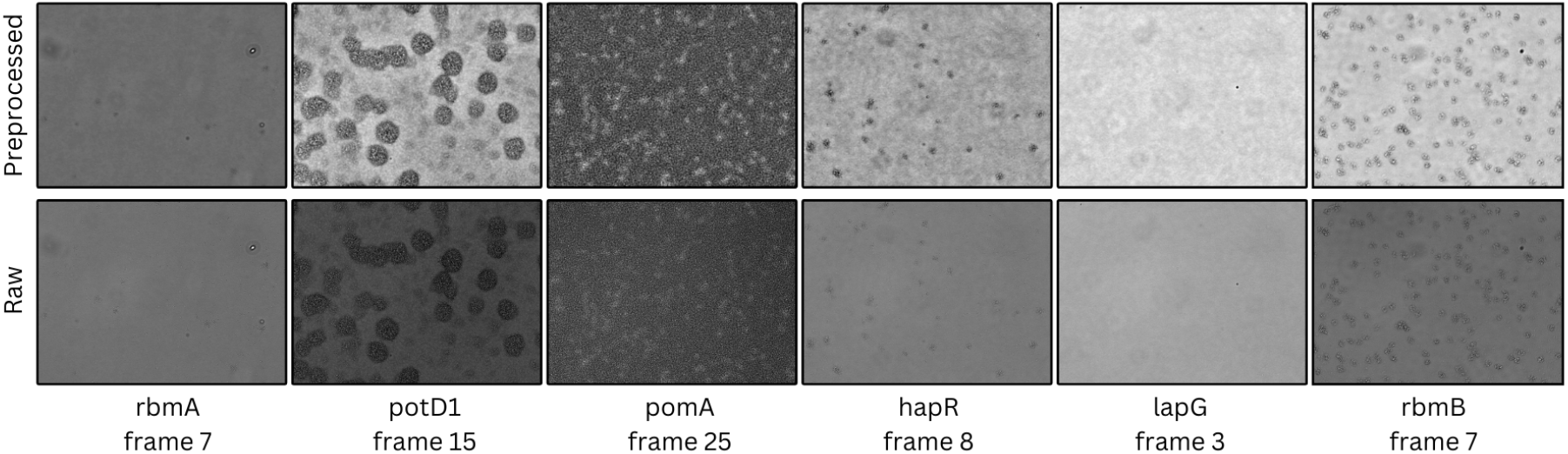
Visualization of frames before (bottom row) and after (top row) preprocessing.

This structure starts with a linear layer that maps input features from 512 (ViT) or 1024 (ResNet50) to 256, followed by a rectified linear unit (ReLU) activation. To add regularization, a dropout layer with a probability of 0.5 is introduced. The subsequent linear layer transforms the output to the desired dimension, and finally, a softmax activation is applied along the second dimension (dim=1) for the final classification.

## 4 Dataset

The dataset used in our research is comprised of 849 videos made up of 10 distinct classes, one class being the parent wildtype (WT) V.cholerae strain, and nine others representing single gene modifications that are known to modify V.cholerae behavior. Briefly, Δ*lapG*, Δ*potD1*, Δ*hapR*, Δ*rbmA, vpvC(w240R)*, Δ*rbmB*, Δ*vpsL* all exhibit unique biofilm characteristics, whereas Δ*flgA* and Δ*pomA* have defects in swimming motility [74, 75]. Each video in the dataset consists of 25 frames, each frame being an image captured hourly for 24 hours, with the first frame being taken at the start. This gives us a longer period of observation. Classes Δ*lapG*, Δ*potD1*, Δ*hapR*, Δ*rbmA* are equally represented, each comprising 96 videos, thereby ensuring a balanced data distribution within these categories. In contrast, the classes Δ*vpsL*, Δ*flgA*, and WT are more densely represented with 126 videos each, offering a substantial dataset for specific analysis in these areas. Conversely, the *vpvC(w240R)*, Δ*pomA*, Δ*rbmB* classes encompass 28, 29, and 30 videos, respectively, ensuring that the dataset includes more specialized subsets. This provides a broad spectrum of data for rigorous analysis and holds potential for significant insights in the field.

**Table 1:**
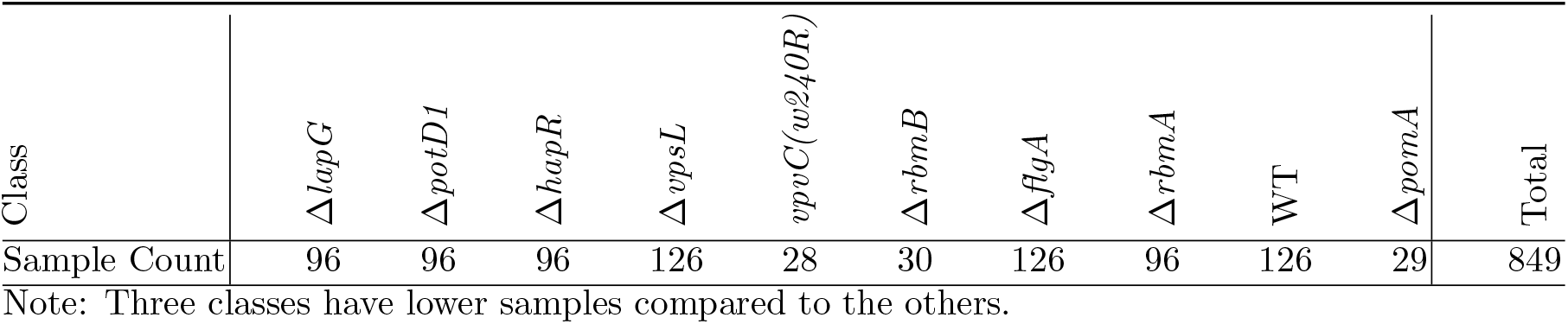
List of classes in the dataset and the number of samples in each class. Three classes have lower samples compared to the others.

| Class | $\Delta lapG$ | $\Delta potD1$ | $\Delta hapR$ | $\Delta vpsL$ | $vpvC(w240R)$ | $\Delta rbmB$ | $\Delta flgA$ | $\Delta rbmA$ | WT | $\Delta pomA$ | Total |
| --- | --- | --- | --- | --- | --- | --- | --- | --- | --- | --- | --- |
| Sample Count | 96 | 96 | 96 | 126 | 28 | 30 | 126 | 96 | 126 | 29 | 849 |

Note: Three classes have lower samples compared to the others.

## 5 Experiments

### 5.1 Visual Inferences

Given the dataset’s substantial volume of high-dimensional video data, we employ dimensionality reduction for ease of analysis. A t-SNE plot visualization of each class’s mid-life point can be observed in Figure 3. We prefer t-SNE as it preserves the local structure of the data, points that are close in the high-dimensional space are likely to be close in the t-SNE plot as well.

**Figure 3:**
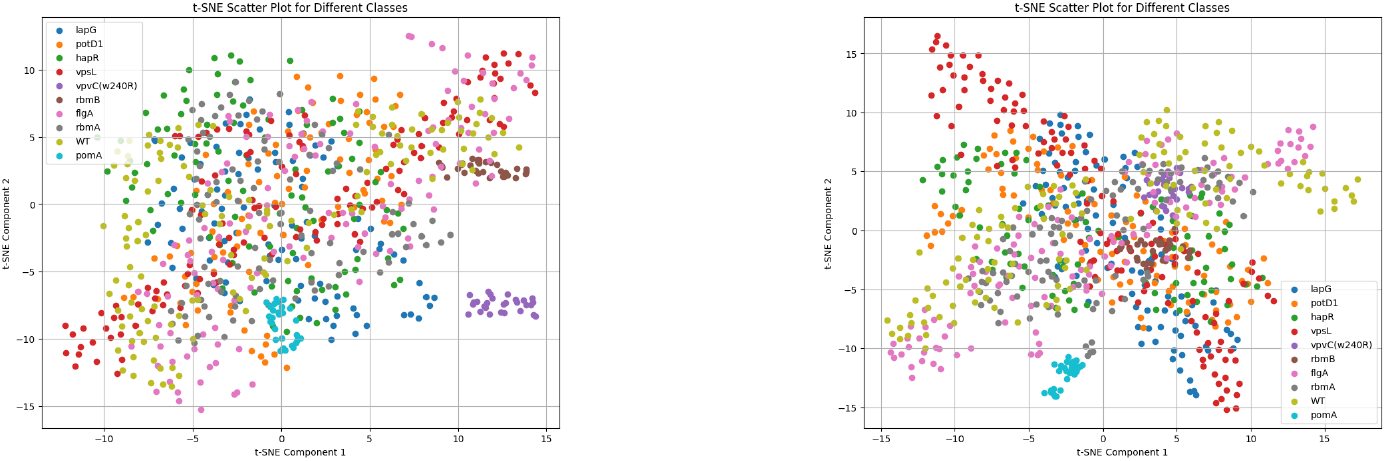
Visualization of the t-SNE graph of halfway life frame features of each class computed for CLIP with ViT-b/32 and ResNet50 backbones, we observe distinct cluster structures for certain classes.

The plot is computed by stacking the mid-life feature frame of each class’s samples and performing dimensionality on this new structure using the TSNE function in sklearn with a perplexity of 20.

We observe that the class Δ*pomA* and Δ*rbmB* form distinct clusters in both ViT-b/32 and ResNet50 backbones of CLIP. Similarly, class *vpvC(w240R)* features are considerably distinct; it is present in the region of Δ*vpsL*, Δ*lapG*, and WT in the case of ResNet50 and is very distinctly clustered in ViT-b/32. We also observe that across both backbones, Δ*vpsL* and Δ*potD1* have a significantly higher spread across the plot.

### 5.2 Implementation Details

We partition the dataset into a train/test split ratio of 75:25 for all conducted experiments. The results are summarized in Table 2.

**Table 2:**
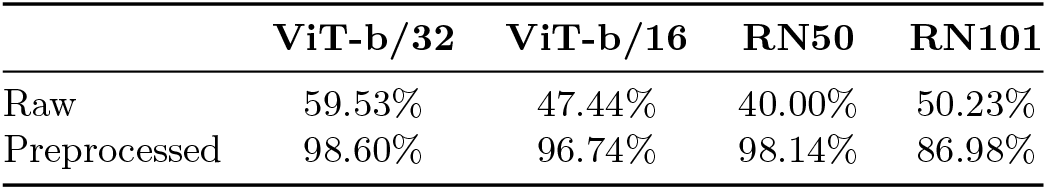
Comparison of the performance of different backbones on raw and preprocessed videos. We observe that preprocessing has a notable impact on the prediction performance.

We employ CLIP backbone with different architectures to extract features for individual frames of each video using pre-trained ViT-B/32, ViT-B/16, ResNet50, and ResNet101 backbones.

The classification of bacteria videos is done using a custom neural network the input dimension is fixed at 512 for ViT-b/32, ViT-b/16, and ResNet101, and 1024 for the ResNet50 backbone which is the dimension of the CLIP embeddings. The Transformer encoder consists of 4 hidden layers, each with a hidden dimension of 512 and 8 attention heads. We train the model with the SGD optimizer set with a momentum of 0.8 and a weight decay of 1*e*^*−*4^, and cross-entropy loss for 1000 epochs with a learning rate of 1*e*^*−*4^ and batch size of 64. Weight initialization using Xavier with a bias of 0.1 is implemented. A learning rate scheduler is also implemented to reduce the learning rate on plateau by a factor of 0.75, and a patience of 50 epochs.

### 5.3 Ablation Study Results

Table 2 presents the classification accuracy of the proposed model using two different backbones on the test set of the bacteria growth dataset. It is noted that although the ViT-b/16 and ResNet101 variants perform well, the best performance is seen with ResNet50 and ViT-b/32.

We observe a notable difference in the performance of predictions between raw data and preprocessed data. Across various backbones, the performance on raw data is comparatively low but increases significantly after preprocessing. This is observed with the ViT-b/32 backbone, the performance jumps from 59.53% on raw data to 98.1% post-preprocessing. Likewise, the ViT-b/16 backbone demonstrates a significant increase from 47.44% on raw data to 96.74% post preprocessing. Moreover, the ResNet50 backbone demonstrates a marked improvement, increasing from 40% on raw data to 98.14% post-preprocessing, and ResNet101 shows an increase from 50.23% on raw data to 86.98% after preprocessing. This shows the effectiveness of preprocessing videos in significantly improving model performance.

### 5.4 Results

The proposed model was tested against the dataset, and we observed the saliency map of each class. Figure 5 presents the saliency maps computed by the ViT-b/32 and the ResNet50 backbones for each class along with the region of standard deviation. The saliency map values are normalized according to min-max normalization to maintain uniformity. We observe that the computed scores are similar, helping us determine what timestamps are of higher interest. Notably for the WT class, which does not exhibit any genetic modifications, we observe that both backbones are nearly identical. However, we also observe that the backbones give us different scores for class Δ*flgA* and Δ*rbmB*, this may be caused due to Δ*rbmB* having lesser training volume.

The proposed model demonstrates notable efficiency in predicting genotypes solely through visual observations of growth patterns captured with low-resolution brightfield microscopy. This highlights the model’s capability to leverage visual cues effectively. In Figure 4, the confusion matrix for the ViT-b/32 and ResNet50 backbones with the proposed model is depicted. We can observe that the overall performance in most classes appears to be excellent for both backbones. Notably, classes such as Δ*flgA*, Δ*vpsL*, and WT have a higher volume of data points, which can be a contributing factor to the model’s robust performance in these categories.

**Figure 4:**
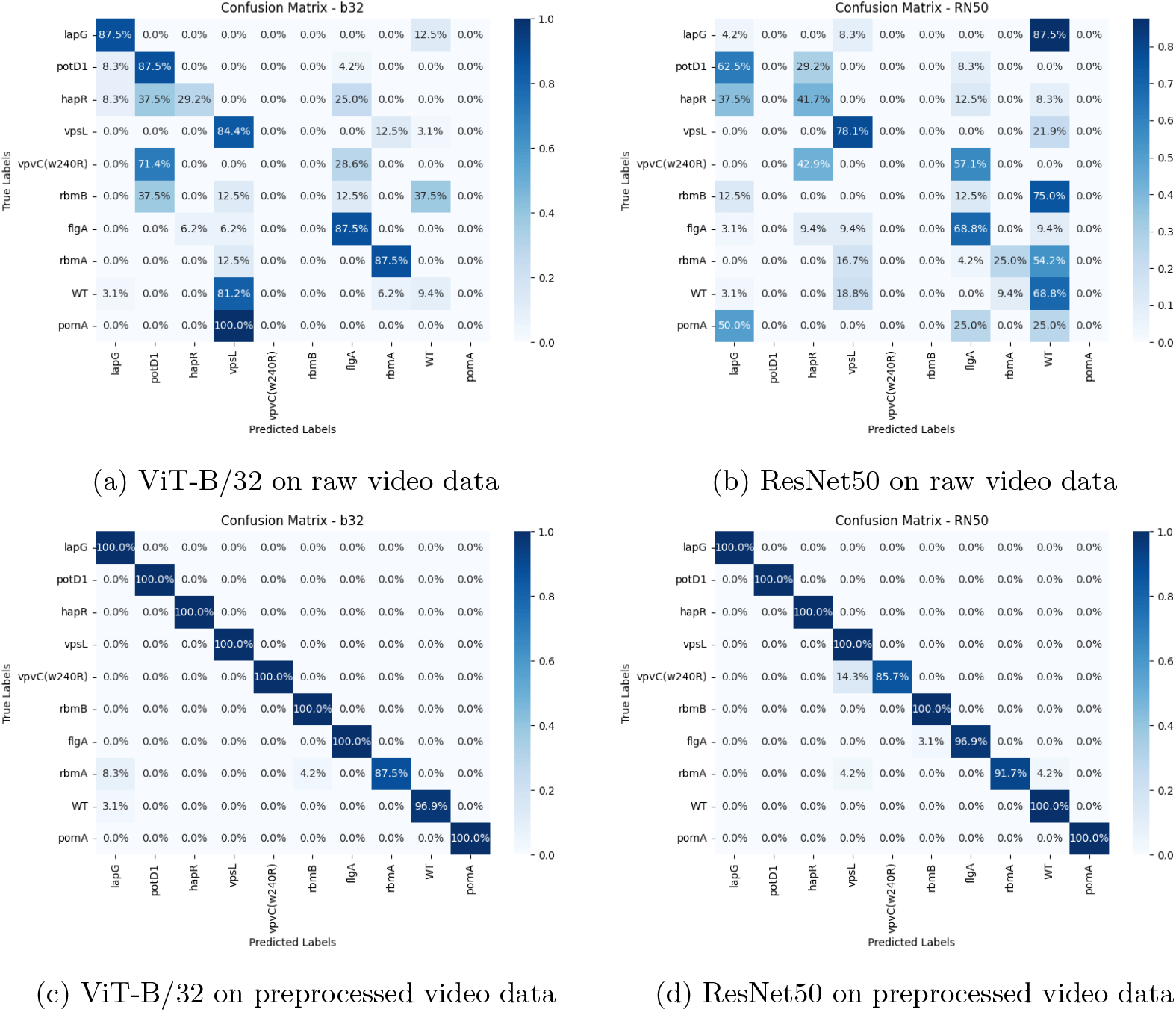
Visualization of Confusion Matrices of the model with features extracted by CLIP using ViT-b/32 and ResNet backbones before (top row) and after preprocessing (bottom row)

**Figure 5:**
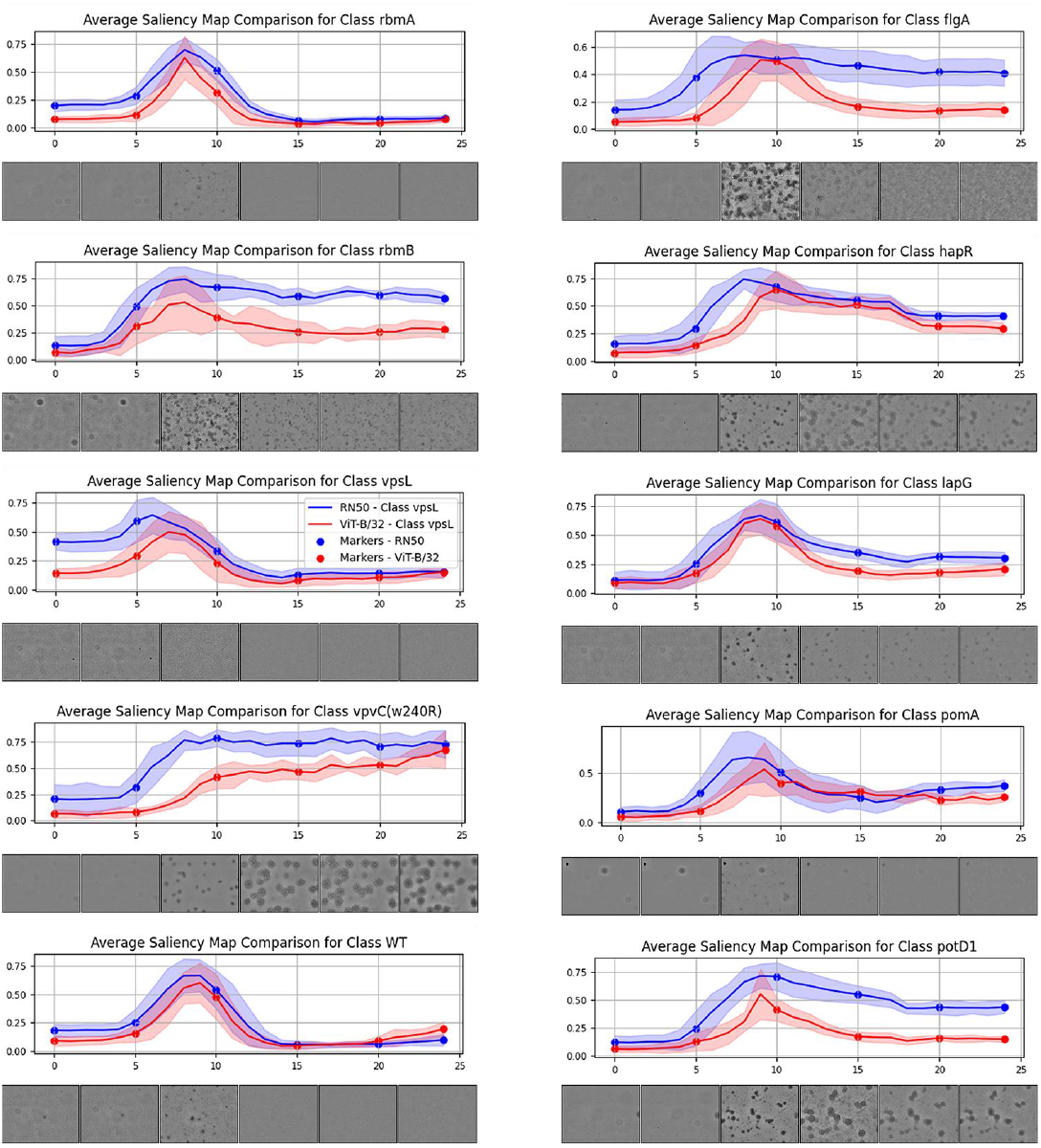
Comparison of average saliency map of each class computed by model with ViT-b/32 (red) and ResNet50 (blue) backbones for CLIP, alongside selected frames at intervals of 1, 5, 10, 15, 20, and 25. We observe a similarity between the graphs of both backbones

However, there are visible misclassifications, predominantly assigning samples to the Δ*rbmA* class in both backbones. Specifically, 8.3% of Δ*lapG* and 4.2% of Δ*rbmB* samples are misclassified as Δ*rbmA* in the ViT-b/32 backbone. Similarly, 4.2% of Δ*vpsL* and WT samples are misclassified as Δ*rbmA* in the RN50 backbone. This suggests that these classes lack distinct visual patterns, leading to confusion for the model. Additionally, we can see very few misclassifications of WT and Δ*flgA*, which can be attributed to the large volume of training data for this class. In the case of *vpvC(w240R)*, it is falsely classified as Δ*vpsL* 14.3% of the time, this falls in line with our observations seen in Figure 3, where *vpvC(w240R)* is seen to have a cluster formed near Δ*vpsL*.

We also observe the superior performance of ViT-b/32 over ViT-b/16 on both raw and preprocessed data. This suggests that the larger vision transformer variant excels in capturing and understanding complex visual features associated with bacteria gene modifications. This highlights the importance of selecting an appropriate backbone for optimal performance.

## 6 Discussion

Our research proposed a supervised deep learning model designed to classify modified bacteria based on known gene modifications explaining the relationship between bacterial phenotype and genotype. Through analysis of bacterial growth behavior captured in videos, we observed a strong correlation between visual patterns and gene modification classes. We introduced a weakly supervised method to identify key moments in bacterial life cycles, thereby enhancing the prediction accuracy. These findings offer valuable insights into how genetic modifications influence bacterial responses, potentially streamlining gene modification research for applications in drug discovery and disease management. Moreover, we can see the potential of predicting genotypes from low-resolution movies that can be acquired with relative ease.

In the goal of refining our model, addressing the observed class imbalance should be a key priority for future enhancements. Classes like Δ*rbmA* have shown instances of misclassification, suggesting that a more balanced dataset is required. Techniques like focused data collection for classes with fewer data points and data augmentation can help to address this imbalance and promote better model generalization. With that in mind, expanding the range of bacterial classes in our dataset could lead to more thorough and precise predictions. This expansion can be achieved by including more bacterial classes in future data collection efforts.

Furthermore, including unsupervised machine learning methods is a promising direction for further exploration. Leveraging clustering or dimensionality reduction methods can reveal underlying patterns in the data, providing insights that improve the predictive capabilities of the model. These approaches collectively can provide a better and more robust model for classifying bacterial gene modification.

## 7 Acknowledgements

This work was supported in part by U.S. NIH grants R01GM134020, P41GM103712, R00AI158939, NSF grants DBI-1949629, DBI-2238093, IIS-2007595, IIS-2211597, and MCB-2205148. This work was supported in part by Oracle Cloud credits and related resources provided by Oracle for Research, and the computational resources support from AMD HPC Fund.

